# Cell-lineage specificity of primary cilia during epididymis post-natal development

**DOI:** 10.1101/288977

**Authors:** Agathe Bernet, Alexandre Bastien, Denis Soulet, Olivia Jerczynski, Christian Roy, Maira Bianchi Rodrigues Alves, Cynthia Lecours, Marie-Ève Tremblay, Janice Bailey, Claude Robert, Clémence Belleannée

## Abstract

Primary cilia are sensory organelles that orchestrate major signaling pathways during organ development and homeostasis. By using a double Arl13b/mCherry-Cetn2/GFP transgenic mouse model, we characterized the spatio-temporal localization of primary cilia in the epididymis, from birth to adulthood. We report here a constitutive localisation of primary cilia in peritubular myoid cells and a dynamic profiling in differentiated epithelial cells throughout post-natal development. While primary cilia are present at the apical pole of the undifferentiated epithelial cells from birth to puberty, they are absent from the apical pole of the epithelium in adults, where they appear exclusively associated with cytokeratin 5-positive basal cells. Exogenous labeling of primary cilia marker Arl13b and IFT88 confirmed the cell lineage specific localization of primary cilia in basal cells and myoid cells in human epididymides. From whole epididymis tissues and serum-free cultures of DC2 murine epididymal principal cell lines we determined that primary cilia from the epididymis are associated with the polycystic kidney disease-related proteins polycystin 1 (PC1) and polycystin 2 (PC2), and Gli3 Hedgehog signaling transcription factor. Thus, our findings unveil the existence of primary cilia sensory organelles, which have the potential to mediate mechano/ chemo-signaling events in the epididymis.

## Introduction

Post-natal development (PND) of the epididymis is a multistep process synchronized with an increase in androgen plasma level, arrival of testicular flow, and sperm release [1]. Proper epididymis PND is essential to generate a fully functional organ that will ensure optimal post-testicular sperm maturation at the time of puberty (For review, [2]). Thus, identification of cellular factors involved in the development of male reproductive functions is a compelling research avenue that may help understand the aetiology of some unexplained male infertility issues.

The adult epididymis is a single convoluted tubule divided into three to four main anatomical regions (*i.e.* initial segment in rodents, *caput*, *corpus* and *cauda*), each displaying distinct morphological features. The lumen of the epididymis through which spermatozoa transit is surrounded by a pseudostratified epithelium. The latter is mainly composed of principal, clear and basal cells, whose functions are essential to ensure proper post-testicular sperm maturation (for review [3]). The epididymis becomes fully functional following a series of morphological changes that occur during its PND, particularly prior to puberty. These changes include the formation and maintenance of a blood-epididymis barrier, the differentiation and organization of epididymal cells to form a well-orchestrated pseudo-stratified epithelium, and the regionalized expression of epididymal genes in the different segments of the organ.

The differentiation of epididymal cell populations and the establishment of distinct epididymal segments was studied in different species during the 80’s, including in mice, rats and humans [1, 4–11]. It consists of three main stages: the undifferentiated period, the period of differentiation, and the period of expansion [12]. During the undifferentiated period (1^st^ week in mouse), the epididymis epithelium is characterized by the presence of columnar cells that lack stereocilia and form primitive junctions. The period of differentiation (2^nd^ to 5^th^ week in mouse) consists in the appearance of differentiated and specialized cells, including clear, principal and basal cells [13, 14]; this period is associated with high cellular proliferation within the initial segment. It has been proposed that the flow of testicular-derived fluid entering the epididymis before sperm production may stimulate this proliferation/differentiation stage [1]. Finally, the period of expansion (6^th^ to 8^th^ week in mouse) features the appearance of spermatozoa within the epididymal lumen, as well as an increase in the size of the epididymis, until cell division arrest at around the 10^th^ week. Different factors such as luminal flow, lumicrine components and androgens have been shown to participate in epididymis PND through epithelial cell differentiation [15], [7, 16]; For review refer to [17]. However, the cellular mechanisms involved in this process are unknown. The serendipitous finding of the presence of sensory primary cilia organelles in the mammalian epididymis shed light on new potential developmental mechanisms [18].

The primary cilium is a solitary cell extension that serves as a sensory organelle and a signaling hub to control cell proliferation, migration, differentiation and planar polarity [19, 20]. Thus, this biological antenna is required to ensure proper tissue development and homeostasis and extends from the surface of most mammalian cells at the post-mitotic stage. Non-motile primary cilia are composed of a 9+0 axoneme (nine pairs of microtubules without central pair, in contrast to 9+2 motile cilia) and a basal body, which derives from the mother centriole of the centrosome. Depending on the biological system in which they are studied, primary cilia play two distinct roles: one as a signaling mediator through a variety of ciliary receptors and their downstream effector molecules, and the other as a mechanosensor through ion channels and transporter proteins. The signaling pathways coordinated by primary cilia are diverse and include Hedgehog (Hh)[21], Wingless (Wnt/Notch) [22] and Platelet-derived growth factor receptor (PDGFR) [23] signaling pathways. In addition to this function, some cells are responsive to the shear stress exerted by the surrounding biological fluids through primary cilia mechanosensing [24], for review [25]. For instance, primary cilia found at the surface of renal intercalated cells, sense urine flow and control cell proliferation through ciliary Polycystin 1 (PC1), transient receptor potential channel interacting, and the Polycystin 2 (PC2) cation channel[26]. Bending of the primary cilium triggered by flow within the nephron tubule induces calcium influx via PC2 calcium transporter, and the downstream regulation of target gene expression involved in cell proliferation [26].

Based on remarkable studies asserting the role of primary cilia in cell sensing, the functional features associated with this organelle make it an ideal candidate for detecting and transducing signals involved in organ development. Apart from the serendipitous observation of primary cilia in the epididymis of equine species reported by Arrighi in 2013 [18], no further studies have investigated the characterization and potential role of these biological antennae in epididymis development and homeostasis. In our study, we discovered for the first time the presence of a primary cilia component in the human epididymis and portrayed their spatio-temporal localization during epididymis PND in a double transgenic mouse model Arl13b-mCherry/Centrin2-GFP [27]. Of relevance, impairment of primary cilia function is associated with male infertility issues and other human diseases referred to as ciliopathies [28]. For instance several reports indicate that Autosomal Polycystic Kidney Disease (APKD), a ciliopathy affecting one in 800-1000 live birth, is associated with a higher prevalence of obstructive azoospermia, increased epididymis and *vas deferens* volumes, and the presence of cysts in the epididymis, seminal vesicles and prostate [29–31]. In addition, Von Hippel-Lindau Syndrome (VHL), an autosomal dominant ciliopathy characterized by the predisposition for multiple tumours, is associated with the development of epididymal cystadenomas that result in male infertility when bilateral [32–35]. Considering that non-genetic environmental insults, such as lithium treatment and folic acid uptake, have been shown to alter cilia length and functions [36–40], portraying and unravelling the role of primary cilia during epididymis PND will open new avenues concerning the diagnosis of unexplained male infertility cases.

## Material and Methods

### Human tissues and ethical consent

Human epididymides were obtained from donors between 26 and 50 years of age through our local organ transplantation program (Transplant Quebec, QC, Canada) after obtaining written consent from the families. Experiments performed in our study were in compliance with the Declaration of Helsinki and were conducted according to the policies for Human Studies with ethical approval from the Centre Hospitalier Universitaire’s (CHUQ) Institutional Review Board (#2018-4043). Human epididymides from three donors were processed as previously described (ref [41]). In brief, the testicles were removed under artificial circulation to preserve tissues assigned for transplantation and processed within 6 h. The epididymides were dissected into three segments, *i.e.* caput, corpus and cauda and fixed by immersion in paraformaldehyde 4% for immunohistological localization of Arl13b and IFT88.

### Mouse tissues and ethics

Epididymides from C57BL/6 and Tg(CAG-Arl13b/mCherry)1Kvand Tg(CAG-EGFP/CETN2)3-4Jgg/KvandJ (referred to as Arl13b-Cetn2 tg in this study, Jackson Laboratory stock# #027967) were used in this study. The Arl13b-Cetn2 tg double transgenic mice express both the ciliary component ADP-ribosylation factor-like protein 13B (Arl13b) fused to the monomeric red fluorescence protein mCherry and the centriolar protein Centrin2 (Cetn2) fused to GFP [27]. These mice were housed and reproduced in the elite animal facility of the CHU de Quebec research Center. Animal experiments were approved by the ethical committee of the Institutional Review Board of the Centre Hospitalier Universitaire de Québec (CHUQ)(CPAC licenses 2016050 and 12016051, C. Belleannée) and were conducted in accordance with the requirements defined by the Guide for the Care and Use of Laboratory Animals.

Epididymides were obtained from mice sacrificed at different post-natal ages. Post-natal day 1 (1 dpn) to 268 dpn tissues were either 1) directly snap frozen in liquid nitrogen and stored at −80°C until use, 2) fixed with fixative containing 4% paraformaldehyde, 10 mM sodium periodate, 75 mM lysine, and 5% sucrose in 0.1 M sodium phosphate buffer (PLP) by immersion (1 to 28 dpn) or by perfusion via the left ventricle (28 to 56 dpn) as previously described [42], or 3) fixed by intra-cardiac systemic fixation with 4% paraformaldehyde for transmission electron microscopy (TEM) analysis as previously described and adapted [43].

### Cell culture

Immortalized Distal Caput principal cells (DC2) developed from mouse tissues were kindly provided by Marie-Claire Orgebin-Crist [44]. DC2 cell were maintained in Iscove’s Modified Dulbecco’s Media (IMDM, Gibco, Invitrogen S.A.) containing 1 μM dihydrotestosterone (Fluka), 10 % of fetal bovine serum (FBS) (Gibco, Invitrogen S.A.) and 50 U ml^−1^ of penicillin G and 50 μg ml^−1^streptomycin (Gibco, Invitrogen S.A.). Cells were kept in an incubator at 32,8 °C in 5% CO_2_ in air and 100% humidity. On the day before the experiment, DC2 cells were plated on fibronectin treated coverslips in a 6-well plate at a density of a 125 000 cells per well. Cells were starved from serum overnight prior to being fixed and used for immunofluorescence staining.

### Immunofluorescence staining

Epididymides from Arl13b-Cetn2 tg mice fixed with PLP were treated for immunofluorescence as previously described [45]. In brief, after three washes in PBS, tissues were cryoprotected with 30% sucrose in PBS for several hours at 4°C, embedded in Tissue-Tek^®^ O.C.T. Compound (Sakura^®^ Finetek, USA), and quick-frozen. Five to 25-μm-thick sections were cut on a cryostat (Shandon Cryotome, Thermo) and collected onto Superfrost/Plus slides (Superfrost Fisherbrand^™^). For indirect immunofluorescence staining, sections were hydrated for 15 min in PBS and treated for 4 min with 1% Sodium Dodecyl Sulfate (SDS) and 2% Triton X-100 in PBS. Sections were washed in PBS for 5 min and then blocked in PBS containing 1% BSA for 15 min. Sections were then incubated overnight in a humid chamber at 4°C with primary antibodies diluted in DAKO solution (DAKO Corp., Carpinteria, CA) and directed against specific markers of basal, principal, clear and myoid cells as well as primary cilia components **(Table 1)**. When specified, an antigen retrieval step for 10 min at 110*°*C in citrate buffer was included prior to incubation with the primary antibodies. Sections were washed in high-salt PBS (2.7% NaCl) twice for 5 min and once in normal PBS. Respective secondary antibodies were then applied for 1 h at room temperature followed by washes, as described above. Characteristics of these antibodies are described in **Table 1**. Slides were mounted in Vectashield medium containing 4’,6’-diamidino-2-phenylindole (DAPI; Vector Laboratories, Inc., Burlingame, CA) for imaging.

**Table 1.**
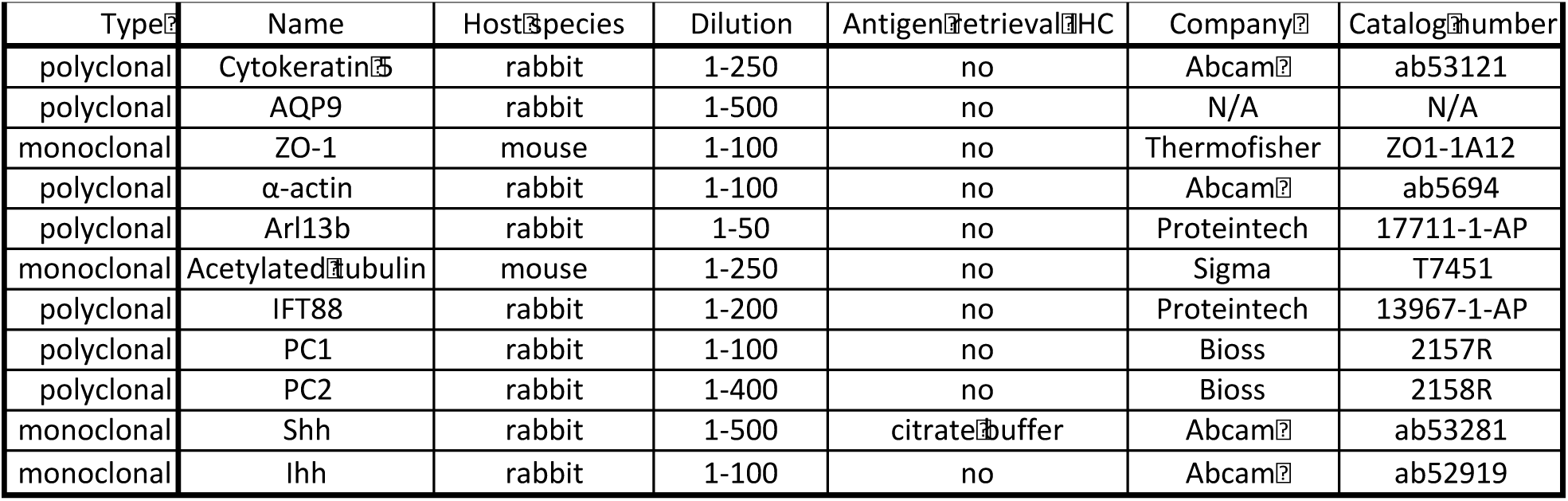
List of antibodies used in our study.

For Immunofluorescence assays performed on cells, the latter were plated on fibronectin coated slides at a density of 125 000 cells per well and fixed with 4% PFA for 10 min. After washes with PBS, blocking was performed for 30 min in PBS solution with 1% BSA and 0.1% Triton X-100. Slides were incubated for 1 hr. with primary antibody (1:400) followed by washes, and for 1 hour with the appropriate secondary antibody (1:800). Slides were mounted in Vectashield medium, as described above.

### Confocal imaging

Digital images were acquired by confocal microscopy on an inverted Olympus IX80 microscope equipped with a WaveFX-Borealin-SC Yokagawa spinning disk (Quorum Technologies; CFI equipment to SE) and an Orca Flash4.0 camera (Hamamatsu). Image acquisition was performed using Metamorph software (Molecular Devices). Optical Z-sections were acquired for each channel and projected into a single picture using ImageJ. Low (20×) and high (100×) magnification pictures were taken for the three major epididymis segments, the *caput*, *corpus* and *cauda* epididymis.

Three-dimensional animations were generated with Bitplane Imaris software v7.5 (Bitplane, Zurich, Switzerland) using images acquired by an Olympus FV-1000 confocal microscope (Olympus Canada; CFI equipment to DS) equipped with a PLAPON60XOSC objective lens (NA 1.4). Isosurfaces were generated in the surpass module, and the camera was rotated to show structures of interest. Generated isosurface renderings were rotated, panned, and zoomed while recording the animation.

To determine primary cilia features throughout the epididymis, images were acquired on a Zeiss LSM700 confocal microscope with a 40x/0.95 plan-apo lens. The resolution was set to the highest possible according to the lens in use (0.145 µm/pixel). Whole epididymal sections (25 µm of thickness) were imaged in the z-axis, generally 15-20 µm at 1 µm interval and large regions of at least 1 mm² were imaged in the XY plane with mosaic tiling at 10% overlap. Excitation was done with lasers at 405, 488 and 555 nm while emission was selected with a variable dichroic splitter. Large data processing was done with custom ImageJ [46] macro and Matlab code. Briefly, images were converted from a single CZI mosaic file to multiple TIFFs. Each tile was then compressed along the z-axis using a maximum intensity projection. Tiles were then stitched together using MIST [47]. Gaussian filtering, thresholding and skeletonizing were performed on the eGFP channel to outline distinct tubules. Centrosomes and cilia were detected using ImageJ’s Find Maxima. The centrosome list was filtered in Matlab to keep only the closest to each cilium and cilia without visible centrosome were excluded. We determined the absolute angle of each cilium with a rotating pattern in order to obtain a maximum at the cilium angle relatively to the closest tubule edge. Matlab and ImageJ macro to process cilia orientation from epididymal confocal acquisitions are available on https://github.com/alexandrebastien/Cilia.

### Immunohistochemistry (IHC)

Fixed epididymides from C57BL/6 mice were treated for IHC as previously described [48]. In brief, tissues were dehydrated, embedded in paraffin and stored at 4°C until use. Paraffin sections (5 μm thickness) were deparaffinised in toluene, hydrated, and then, where specified, treated for antigen retrieval with citrate buffer (pH 6.0). Endogenous peroxidase activity was then quenched with 3% H_2_O_2_ (v/v) in methanol for 10 min. Sections were washed for 5 min in 95% ethanol and 5 min in PBS. Non-specific binding sites were blocked with 0.5% BSA for 1 h. Primary antibodies diluted in DAKO were applied overnight at 4°C (**Table 1**). For control sections, PBS replaced the primary antibodies. Sections were subsequently incubated with biotinylated 1) donkey anti-goat or 2) goat anti-rabbit antibodies for 60 min, and with ABC elite reagent (Vector Laboratories, Inc. Burlingame, Ca) for 30 min. Immunostaining was revealed using 3-amino-9-ethylcarbazole (AEC). Mayer’s hematoxylin solution was used for counterstaining, and mounted under cover slips using an aqueous mounting medium (Sigma). Slides were observed under a Zeiss Axioskop 2 Plus microscope linked to a digital camera from Qimaging. Images were captured using the QCapture Pro (Qimaging Instruments).

### Transmission Electron microscopy

#### Tissue Preparation

Mice were anesthetized with ketamine (intraperitoneally) and tissues were fixed by intracardiac perfusion with PBS for 2 min followed by 4% PFA for 2 min and treated as previously described and adapted [43]. In brief, tissues were further fixed by immersion for 2 h in 3.5% acrolein, followed by 3 h in 4% PFA and washed three times in PBS for 15 min. Fifty-micrometer-thick transverse sections of the epididymis were cut in PBS using a vibratome (Leica VT100S) and stored at −20°C in cryoprotectant until further processing.

#### Electron microscopy

Sections stored in cryoprotectant were rinsed in PBS, postfixed in 1% osmium tetroxide for 30 min at room temperature and dehydrated in ascending concentrations of ethanol (from 35% to 100%). Sections were then treated with propylene oxide and impregnated in Durcupan resin (EMS) overnight at room temperature. After mounting between ACLAR embedding films (EMS), the resin was polymerized at 55°C for 72 h. Areas of interest were excised from the embedding films, re-embedded at the tip of resin blocks, and cut at 65–80 nm of thickness using an ultramicrotome (Leica Ultracut UC7). Ultrathin sections were collected on square mesh grids (EMS) and examined at 80 kV with an FEI Tecnai Spirit G2 transmission electron microscope. Pictures were acquired using an Orca-HR camera (10 MP; Hamamatsu).

## Results

### Primary cilia extend from basal and smooth muscle cells in the epididymis of adult mice

Due to the fact that primary cilia are small, dynamic and present as a solitary figure on the cell surface, these organelles are difficult to detect within *in situ* or *in vivo* whole organ systems. We used the Arl13b-Cetn2 tg mouse model [27] to accurately study lineage specificity of primary cilia from whole epididymis tissues. Arl13b-positive structures were observed within the epithelium and in the surrounding smooth muscle layer of the epididymis **(Fig. 1A)**. The microtubule component and primary cilia marker acetylated tubulin (Ac-Tub) was observed associated with the sperm flagellum and co-localized with Arl13b-positive cell extension, further confirming the specific detection of non motile primary cilia from Arl13b-Cetn2 tg mice **(Fig. 1BC)**. By using specific markers of distinct epididymis epithelial cell populations (*i.e.* keratin 5 for basal cells, aquaporin 9 for principal cells and alpha actin for smooth muscle cells) we determined the cell-specific localization of primary cilia from adult Arl13b-Cetn2 tg mice **(Fig. 2).** As evidenced by labelling with the alpha actin smooth muscle cell marker, primary cilia were found in the peritubular area associated with smooth muscle cell centriole **(Fig. 2A, inset).** The ultrastructure of primary cilia in smooth muscle cells is composed of a basal centriole and an axoneme extension as observed by transmission electron microscopy **(Fig. 2B).** In addition, Arl13b-positive primary cilia were never observed associated with aquaporin 9-positive principal cells, nor aquaporin 9-negative clear cells from adult mice **(Fig. 2C).** Finally, we found that primary cilia extended from the centriole of keratin 5-positive basal cells **(Fig. 2D, inset).** From the whole image acquisitions performed on adult epididymis tissues, all basal cells that were observed within a 25 µm tissue section exposed a primary cilium. While the major part of the basal cell body is located at the base of the epithelium, basal cells dynamically extend towards the lumen of the epididymis through axiopodia [49, 50]. According to basal cell 3D reconstruction from transgenic epididymal tissues, we observed that primary cilia take their origin from the centriole located at the apical pole of the cell body and at the base of the axopodia **(Fig. 3)**. In addition, basal cell-associated primary cilia never cross epididymis tight junctions as evidenced by 3D reconstruction after ZO-1 staining **(Supplementary file 1).**

**Fig. 1.**
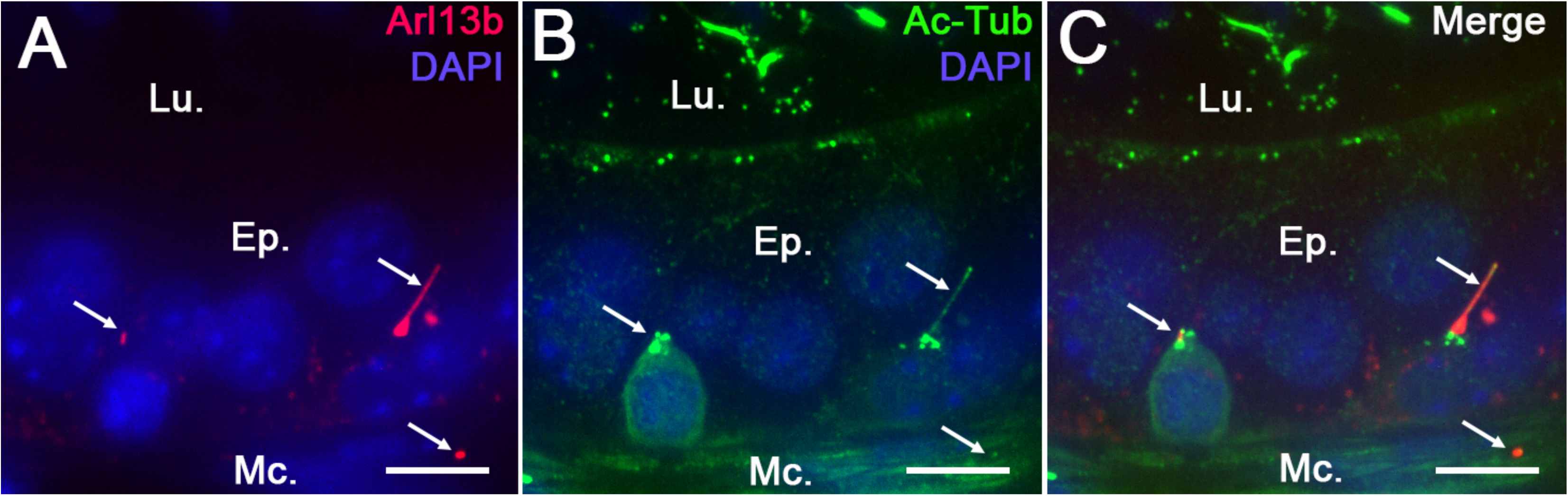
Detection of primary cilia in Arl13b-Cetn2 tg mice. Epididymal sections from Arl13b-Cetn2 tg mice **(A)** were stained for the primary cilia marker acetylated tubulin (Ac-Tub) **(B)**. Arl13b-positive cilia extensions colocalize with Ac-Tub **(Arrows, A, B and C)**, which validates this mouse model for the study of primary cilia in the epididymis. Ep.: epithelium; Lu.: lumen; Mc.: myoid cells. Scale bar: 10 µm.

**Fig. 2.**
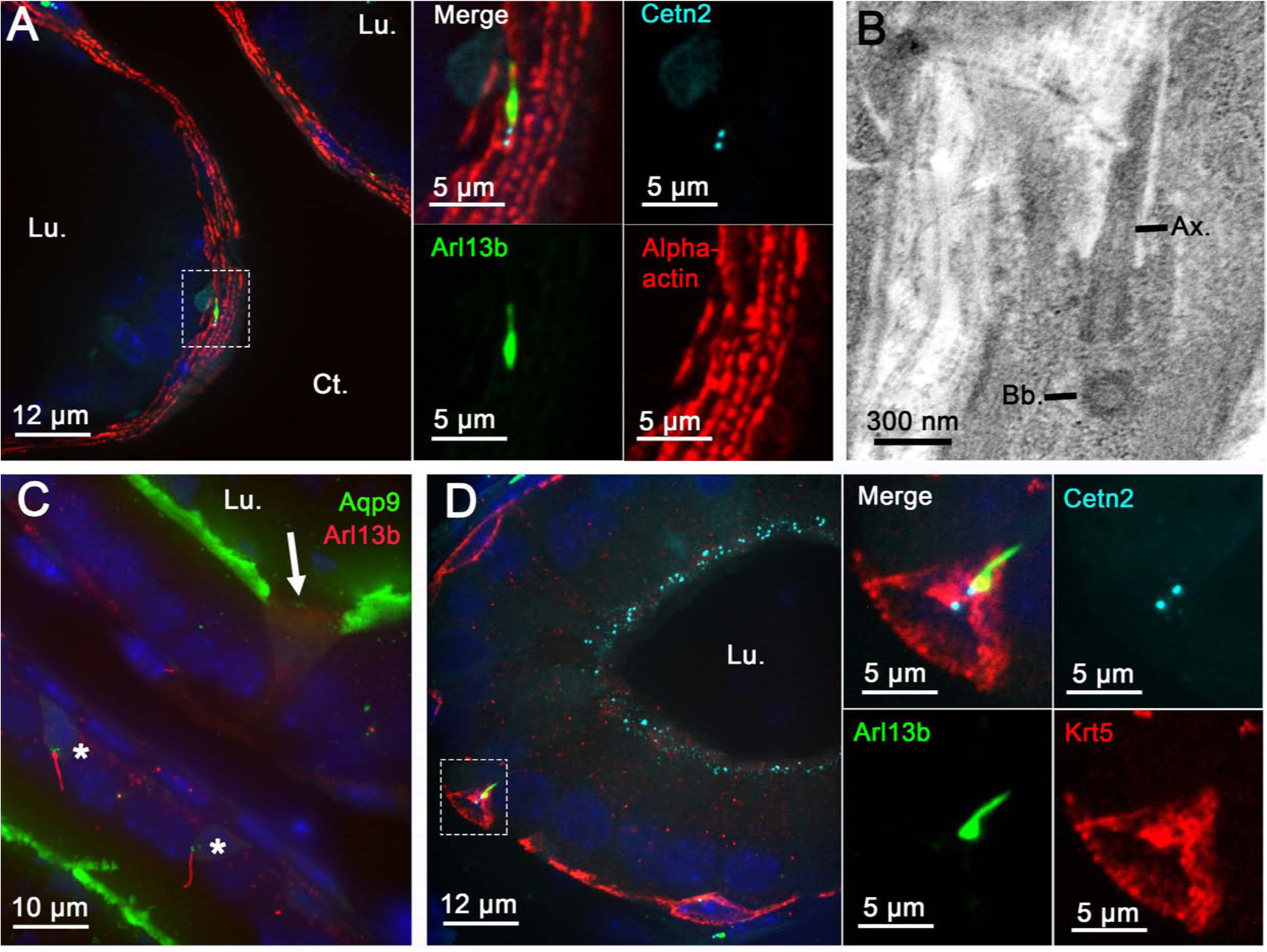
Primary cilia extend from basal and peritubular myoid cells in the epididymis of adult mice. Immunofluorescent staining for alpha-actin myoid cell marker **(A and inset)**, aquaporin 9 (Aqp9) principal cell marker **(C),** keratin-5 (Krt5) basal cell marker **(D and inset)** was performed on epididymal sections from Arl13b-Cetn2 dTg mice. Primary cilia structural components were detected by transmission electron microscopy in epididymal peritubular myoid cells from C57BL/6 mice **(B).** Ax.: axoneme; Bb.: basal body; Ct.: connective tissue; Lu.: lumen; Mc.: myoid cells.

**Fig. 3.**
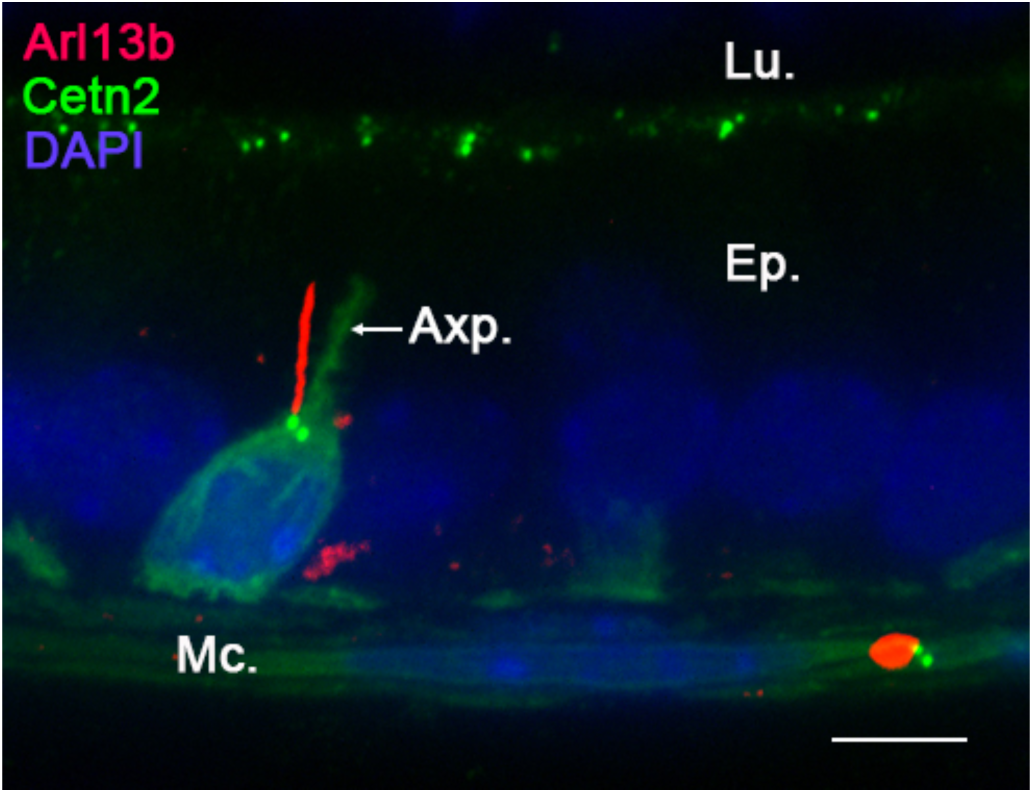
Primary cilia extend from basal cells’ main body. Confocal acquisition of the initial segment from the epididymis of Arl13b-Cetn2 dTg mice showing a ciliated basal cell. Arl13b-positive primary cilia extend from centrin2-positive centrioles located at the base of the axiopodia (Axp.).

### Primary cilia profiling along the adult epididymis

The epididymis is a highly segmented tubule with each segment displaying distinct features in terms of cell populations, cell composition and cell functions [51]. We investigated the properties of basal cell primary cilia throughout the different segments of the epididymis from Arl13b-Cetn2 tg mice (*i.e.* initial segment (IS)*, caput, corpus* and *cauda*) by whole tissue confocal imaging and digital image processing **(Fig. 4)**. Primary cilia were observed associated with peritubular myoid and basal cells in all segments of the epididymis, with a higher proportion in the basal cells from the distal regions **(Fig. 4A, Supplementary file 2)**. In the IS basal cell primary cilia appeared as short 2–3 µm extensions directing towards the base of the epithelium, according to GFP-positive centriole localization **(Figs. 4A, B)**. In the other segments, *i.e* the *caput, corpus* and *cauda*, primary cilia appeared as longer 5–10 µm extensions, presenting with erratic directions. Furthermore, myoid cell primary cilia presented no obvious variation from one segment to another in terms of length (5–6 µm) or orientation (generally perpendicular to the intratubular axis). In addition to their localization throughout the basal and peritubular myoid cells of the epididymis, primary cilia were also associated with basal cells in the *vas deferens* (VD) **(Fig. 4A, VD)** and were observed as tremendously long extensions in the *efferent ducts* (ED) **(Fig. 4A, ED)**. In ED, ciliated cells presenting with numerous centrin2-positive centrioles were observed from the apical surface of the ducts. Moreover, elongated Arl13b-positive primary cilia (15–40 μm) were found associated with centrioles located in non-ciliated cells of the ED.

**Fig. 4.**
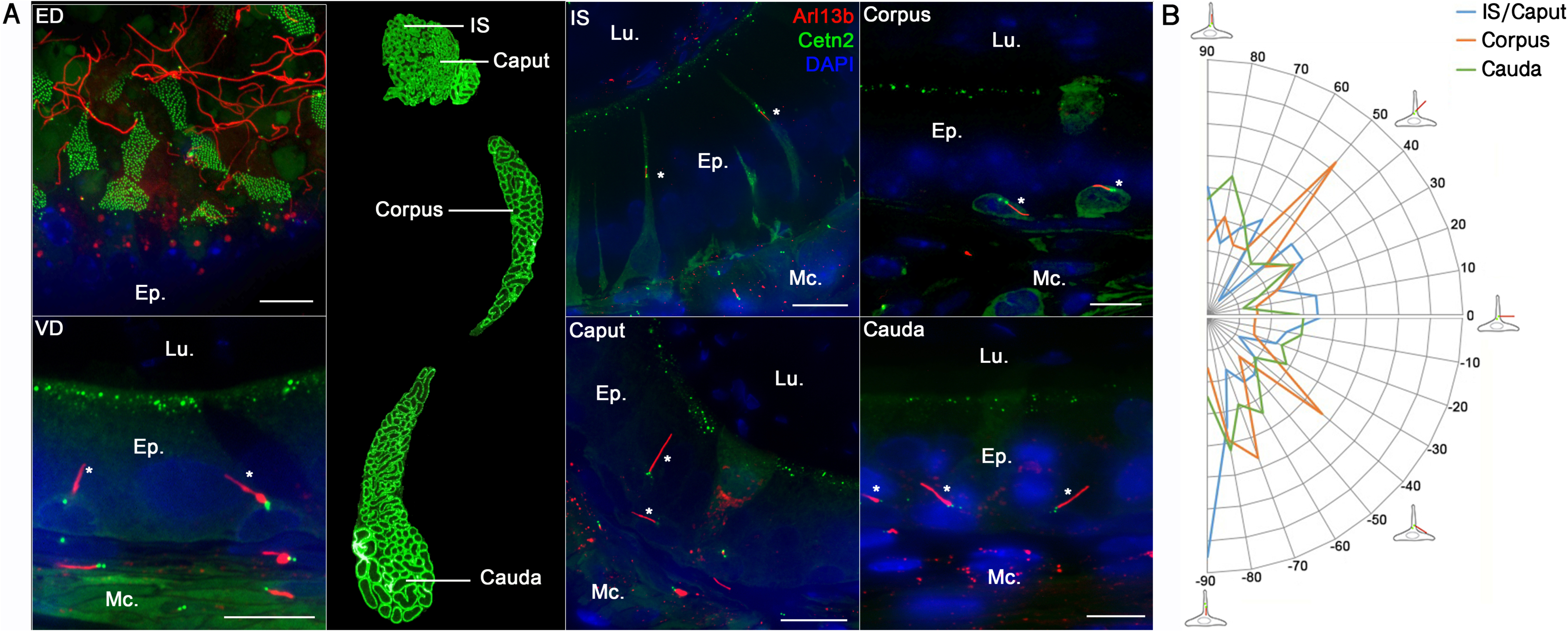
Primary cilia are observed in the efferent ducts (ED), the *vas deferens* (VD), and the different segments of the mouse epididymis. Primary cilia were observed associated with basal cells (stars) and peritubular myoid cells in all segments of the epididymis **(A)**. The angle of basal cell primary cilia relative to the closest tubule edge was determined by image processing in each epididymal region, *i.e.* IS/*Caput* (n=899 cells), *corpus* (n=371 cells) and *cauda* (n=1065 cells) from 3 mice. Proportions of primary cilia with distinct angles were distributed between −90° for cilia directed towards the base of the epithelium to 90 ° for cilia directed towards the lumen **(B)**. Ep.: epithelium. Lu.: Lumen. Mc.: myoid cell. IS: initial segment. Scale bars: 10 µm.

### Dynamic primary cilia exposure during epididymis post-natal development

The epithelium of the epididymis undergoes important changes during the different PND stages [1]. We explored the dynamics of primary cilia exposure that accompanies epididymis epithelium development just after birth (at 5 dpn), at pre- and post-pubertal ages (at 30 and 42 dpn, respectively), and at the adult stage (at 42 to 268 dpn) **(Fig. 5).** The Arl13b-Cetn2 tg mouse model displays the same pattern of epididymis development as previously described in wild-type mice [12] **(Fig. 5, Top).** For instance, H & E staining performed on the epididymis at 5 dpn stage shows undifferentiated columnar cells of the epididymal epithelium surrounding an empty lumen. At 30 dpn, we observed a pseudo-stratified epithelium and the presence of basal cells. The lumen is filled with non-cellular material. At 42 dpn and 268 dpn, the pseudostratified epithelium surrounds large lumens filled with spermatozoa. At an early stage of dpn **(Fig. 5, at 5 dpn Bottom and supplementary file 3)**, primary cilia are observed by confocal microscopy as short Arl13b-positive extensions associated with centrioles, located at the apical surface of non-differentiated columnar cells. Primary cilia extensions are extensively observed associated with myoid cells surrounding the epithelium. At 30 dpn, prior to puberty, primary cilia are found at the apical surface of the epithelium associated with differentiated cells, potential clear and/or principal cells, and associated with basal cells. After puberty, at 42 dpn **(Fig. 5, at 42 dpn and supplementary file 4)** and 268 dpn, primary cilia extensions are exclusively found in basal cells within the epithelium as well as in surrounding myoid cells. Thus, according to the PND stage of the mouse epididymis, primary cilia are localized at different positions along the epididymal epithelium, *i.e.* facing the lumen during the undifferentiated period, facing the lumen and at the base of the epithelium during the period of differentiation, and exclusively at the base of the epithelium during the expansion phase. Primary cilia are observed in myoid cells at all PND stages studied. According to ZO-1 immunostaining of the apical pole of the epididymis **(Fig. 6)**, primary cilia extend from the epithelial cell surface at an early stage of PND (at 5 dpn) and are in direct contact with the intraluminal compartment.

**Fig. 5.**
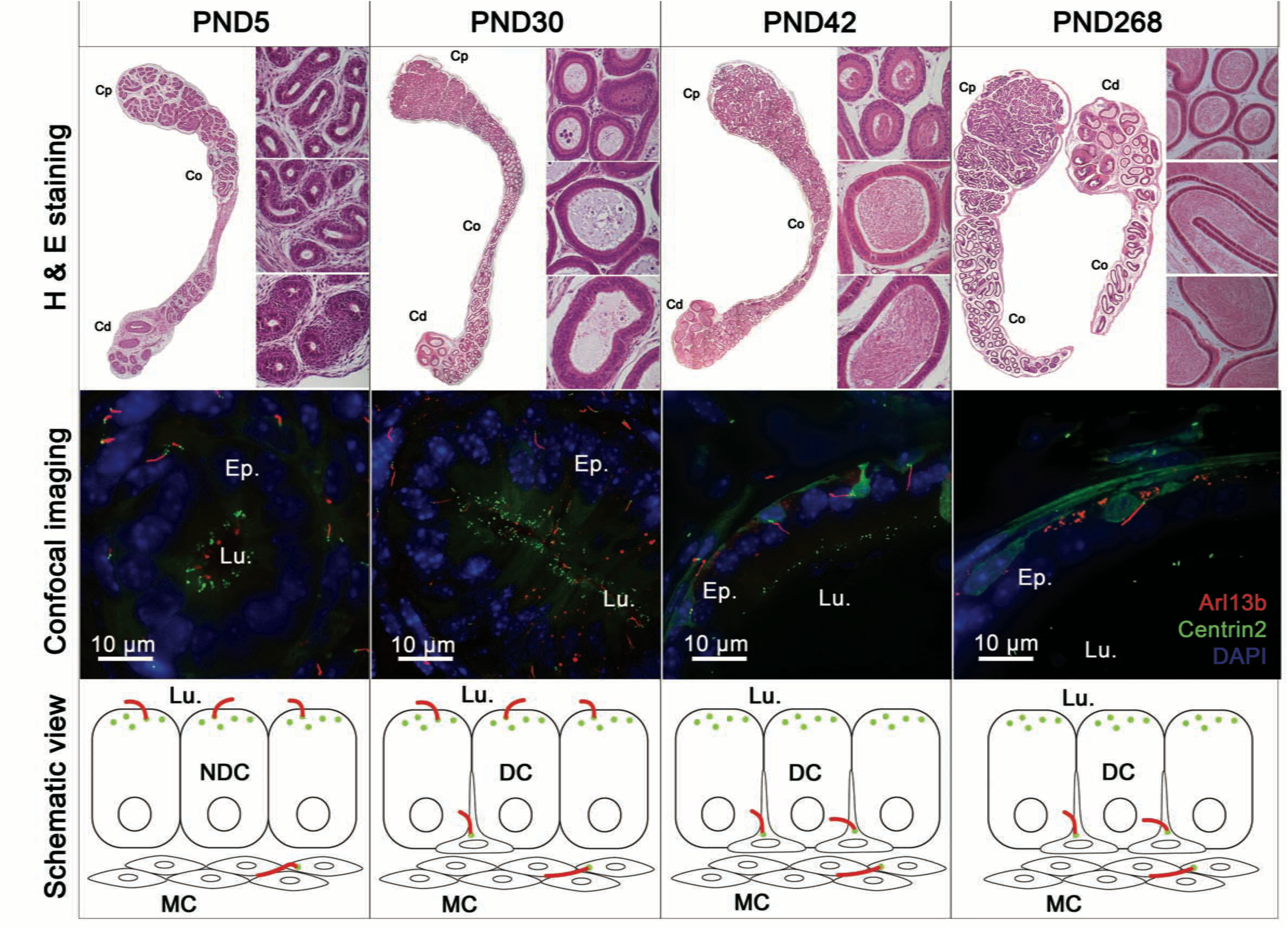
Primary cilia are observed at different stages of epididymis post-natal development. Hematoxylin and eosin staining (H & E) and confocal imaging were performed on epididymis sections from Arl13b-Cetn2 dTg mice at different stages of post-natal development, *i.e.* at 5, 30, 42 and 268 dpn. Schematic representation of primary cilia location is shown throughout post-natal development with centrioles displayed in green and primary cilia in red. NDC: non-differentiated cells; DC: differentiated cells; MC: myoid cells; Lu.: lumen.

**Fig. 6.**
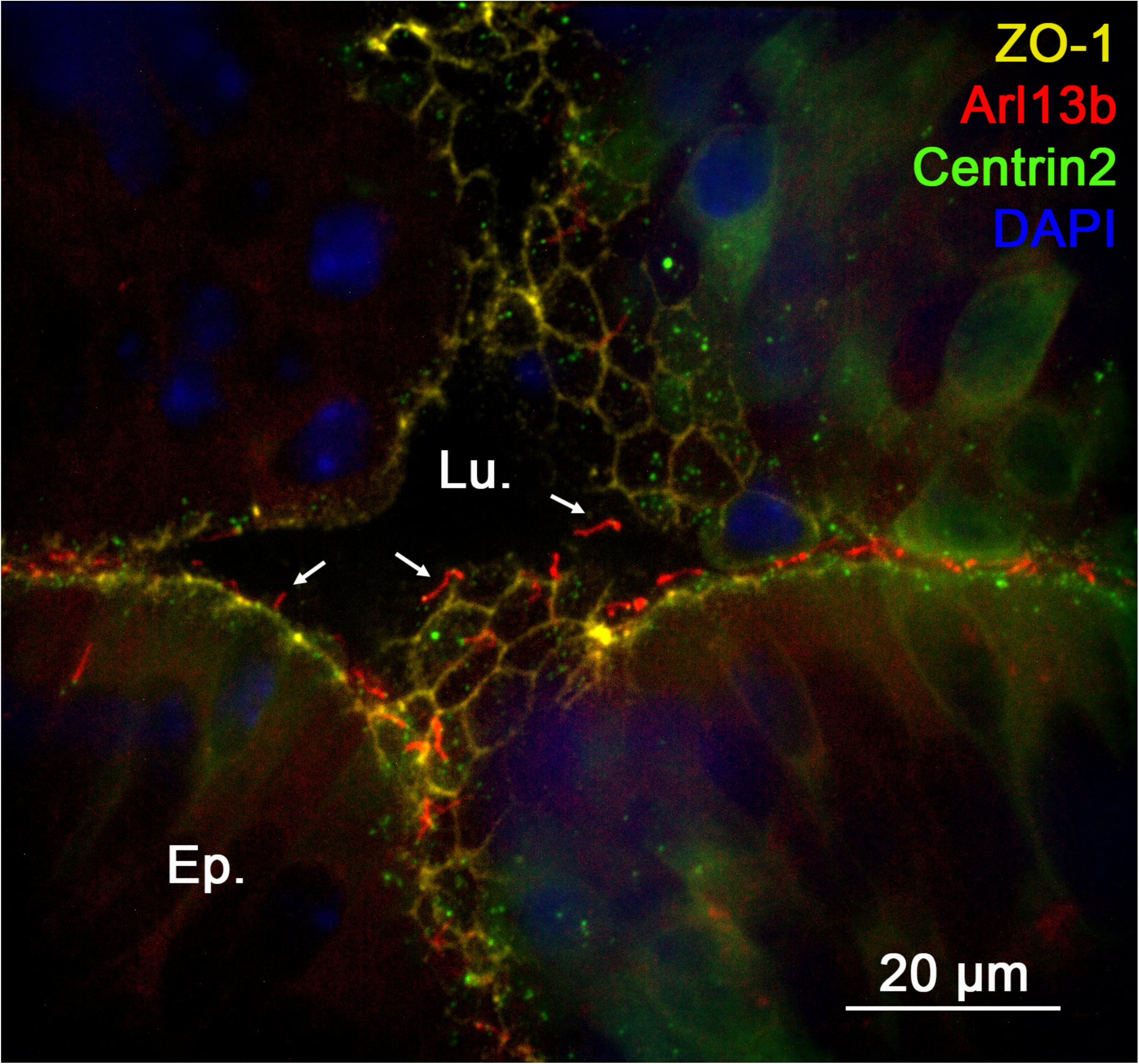
Primary cilia are in contact with the intraluminal compartment of the epididymis at an early stage of post-natal development. Immunofluorescent staining for the tight junction protein-1 (ZO1) on the cauda epididymidis of Arl13b-Cetn2 dTg mice at 5 dpn. Lu.:lumen; Ep.:epithelium.

### Primary cilia of the epididymis are potential mechano/chemo-sensors of the cell-surrounding environment

According to the literature that describes the function of primary cilia in different model systems, two major roles are associated with these biological antennae: the first as a chemo-sensor of major signaling pathways (*e.g.* Hg, Wnt, PDGFR), and the second as a mechano-sensor responsive to shear-stress and pressure exerted on the cell surface by extracellular bodily fluid. By analogy with other tissues, we investigated the functional signaling components associated with primary cilia in the epididymis.

We first explored the expression level of PC1 and PC2 **(Fig. 7)**, two major players in the primary cilia-dependent fluid-flow mechanosensation. At dpn 7, PC1 was detected at the apical pole of undifferentiated cells as well as in peritubular myoid cells in the different epididymal segments **(Fig. 7A).** At this post-natal stage, a stronger signal was observed in the *vas deferens* **(Fig. 7A, VD)**. In tissues from adult mice, PC1 was intensively detected in the cytoplasm and the apical pole of the epithelium in the initial segment/*caput* of the epididymis. While PC1 intensity transitionally decreased in the *corpus* compared to the *caput*, it drastically increased in the apical membrane of epithelial cells from the *cauda* as well as in the *vas deferens* (VD). PC1 was also detected, but to a lower extent, in the peritubular myoid cells. While PC2 could not be detected on epididymal sections by IHC under our experimental conditions (not shown), we observed a co-localisation between PC2 and the primary cilia marker Ac-Tub in the DC2 epididymal principal cell line **(Fig.7B, arrows).**

**Fig. 7.**
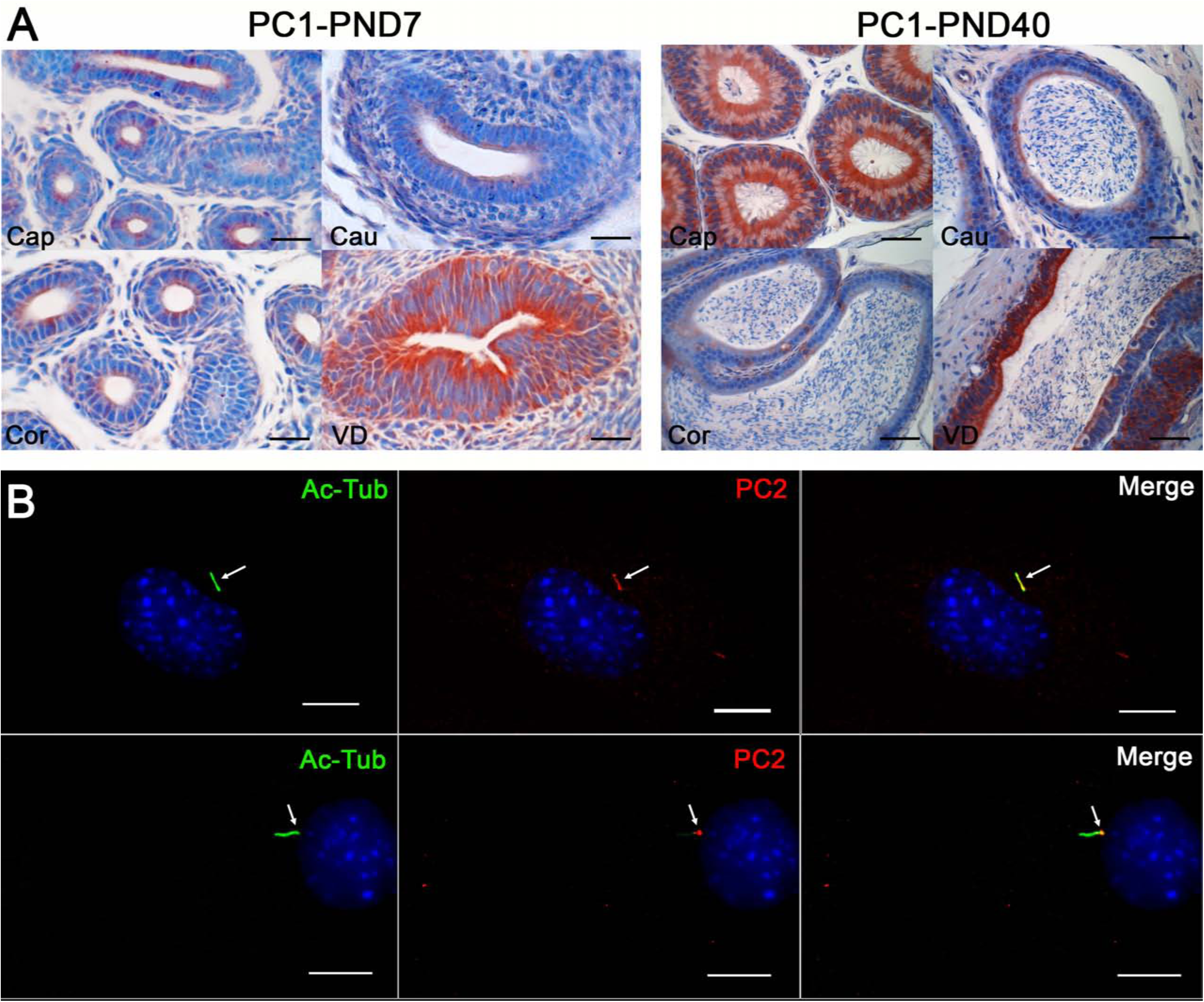
Expression of Polycystin1 (PC1) and Polycystin 2 (PC2) in epididymal cells and in the mouse epididymis. **(A)** Immunohistochemical staining for PC1 on the epididymis of Arl13b-Cetn2 dTg mice at 7 and 40 dpn. Cap.: caput; Cor.:corpus; Cau.:cauda; VD: *Vas deferens*. Scale bar: 50 µm. **(B)** Immunofluorescent staining for PC2 and the primary cilia marker acetylated tubulin (Ac-Tub) in DC2 cells.

We next investigated the expression of GLI3, a downstream transcriptional factor of the Hh pathway **(Fig. 8)**. In adult epididymis, GLI3 was observed as discontinuous staining at the apical pole of the epithelium in the *corpus* and *cauda* epididymidis. In addition, strong GLI3 staining was observed in the nucleus of different epithelial cell populations in all segments of the organ. This staining was particularly intense in basal cell nuclei **(Fig. 8, stars and insets).**

**Fig. 8.**
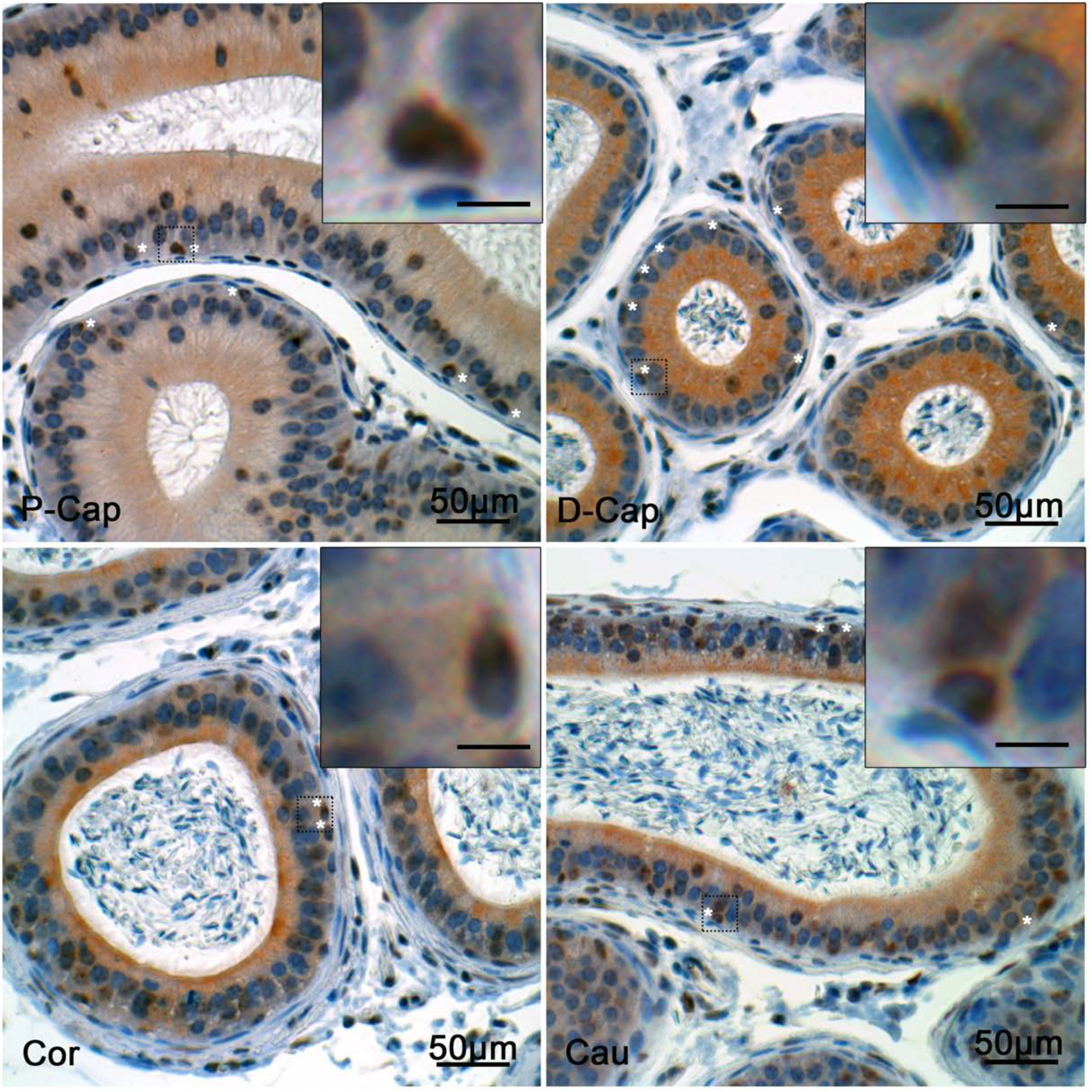
Expression of Gli3 in the mouse epididymis. Immunohistochemical staining for Gli3 on the epididymis of Arl13b-Cetn2 dTg mice at 40 dpn. P-Cap: proximal caput; D-Cap: distal caput; Cor.:corpus; Cau.:cauda. Inset scale bar: 10 µm

### Presence of primary cilia components in the human epididymis

In order to translate our observations made from a transgenic mouse model, we investigated the expression of two primary cilia markers, Arl13b and IFT88, in human epididymal tissues **(Fig. 9, Supplementary file 5).** We observed strong exogenous staining for Arl13b in the apical pole of the epithelial cells from the proximal segment, corresponding to the *efferent ducts/caput* **(Fig. 9, ED/caput inset)**. Whereas the signal was less intense in the *corpus* and *cauda* epididymidis, it was mainly localized in peritubular myoid cells as well as in basal cells **(Fig. 9, arrows in insets)**. An identical detection profile was observed for IFT88 (**Supplementary file 5).**

**Fig. 9.**
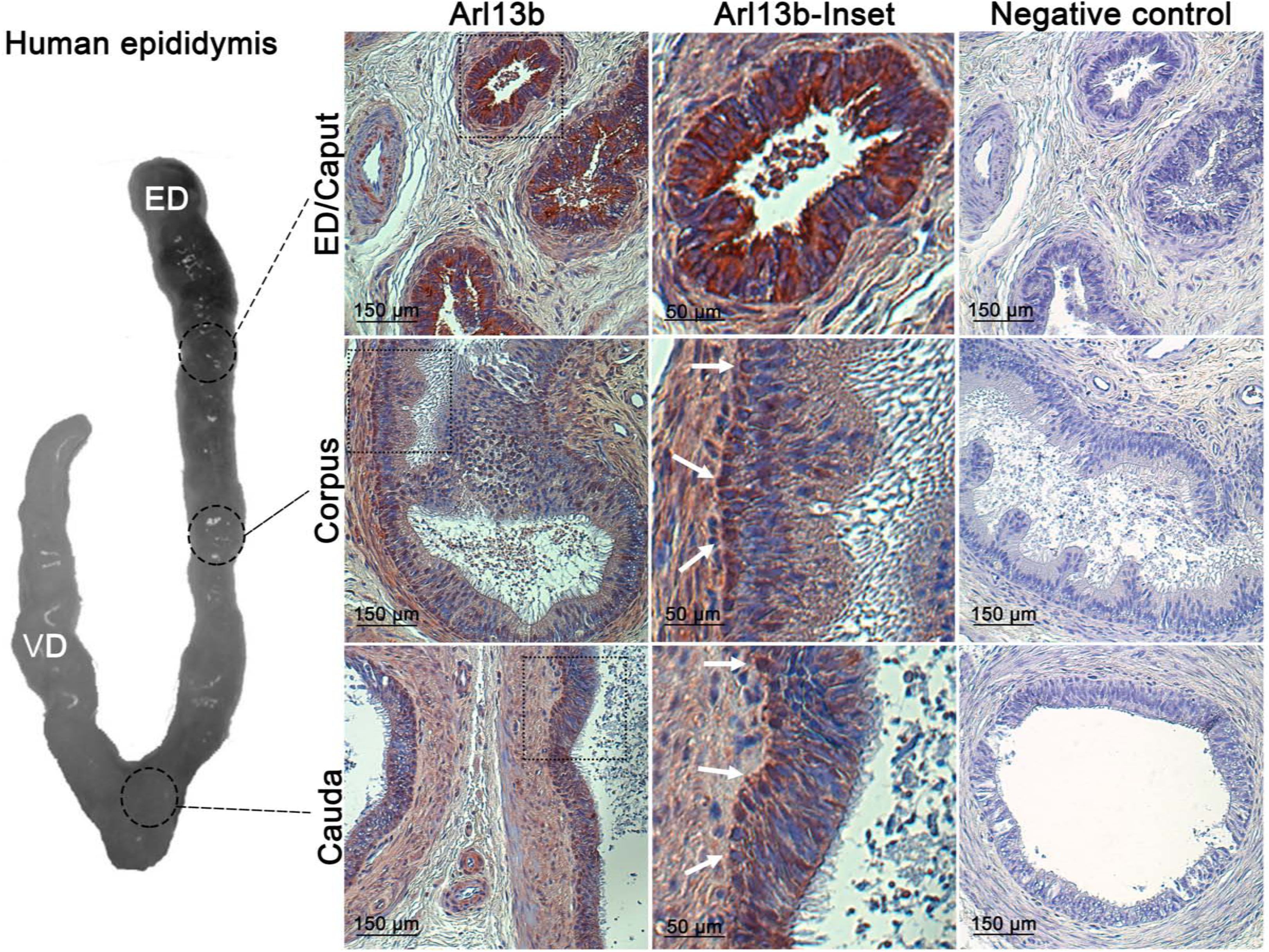
Expression of Arl13b in the human epididymis. Immunohistochemical staining for Arl13b was performed on the different segments of a human epididymis (donor age = 50).

## Discussion

Primary cilia are solitary antennae found at the cell surface and are fundamental to organ development and homeostasis [20]. Accordingly, impairment of primary cilia formation in humans is responsible for a broad range of clinical syndromes—including male infertility—and diseases referred to as ciliopathies [28–35]. While these organelles have been broadly studied in most biological systems, investigations performed on primary cilia in the male reproductive system remain scarce [18, 52]. Acknowledging the potential of primary cilia in the diagnosis of unexplained male infertility, we portrayed for the first time the cell lineage specificity of primary cilia in the epididymis, the organ of the male reproductive system in charge of post-testicular sperm maturation (*i.e.* acquisition of motility and abilities to recognize and fertilize an egg) and sustaining male fertility.

Almost every vertebrate cell has a primary cilium, which is a dynamic cell surface projection playing the role of a sensory and signaling organelle. Although primary cilia were first described by Ecker in 1844, knowledge of the full scope of their functions is continuously expanding with the development of transgenic mouse models and the increasing number of human diseases related to primary cilia dysfunctions [53]. With the re-appreciation of this biological antenna over the past 10 years, it has become clear that the role of primary cilia in sensing the extracellular environment potentially takes place in almost every cell responsive to intercellular cross-talks and bodily fluid changes. The epididymis is a singular tubule that comprises distinct segments, each controlling the composition of a dynamic epididymal fluid for proper sperm maturation. Several mechanisms of intercellular communication between epididymal somatic cells and between somatic and spermatic cells have been shown to be of major importance for the control of sperm fertilizing abilities and reproductive outcomes [45, 50, 54–56]. We unveiled here, for the first time, the existence of primary cilia displaying a cell lineage specificity and key features of signaling antenna in the developing epididymis. In light of the spatio-temporal changes of primary cilia observed during PND, and the functional components associated with these antenna (*i.e.* PC1/PC2 and Gli3), we propose a novel mechanism of intercellular communication involving primary cilia organelles in the control of epididymis development and homeostasis **(Fig. 10).**

**Fig. 10.**
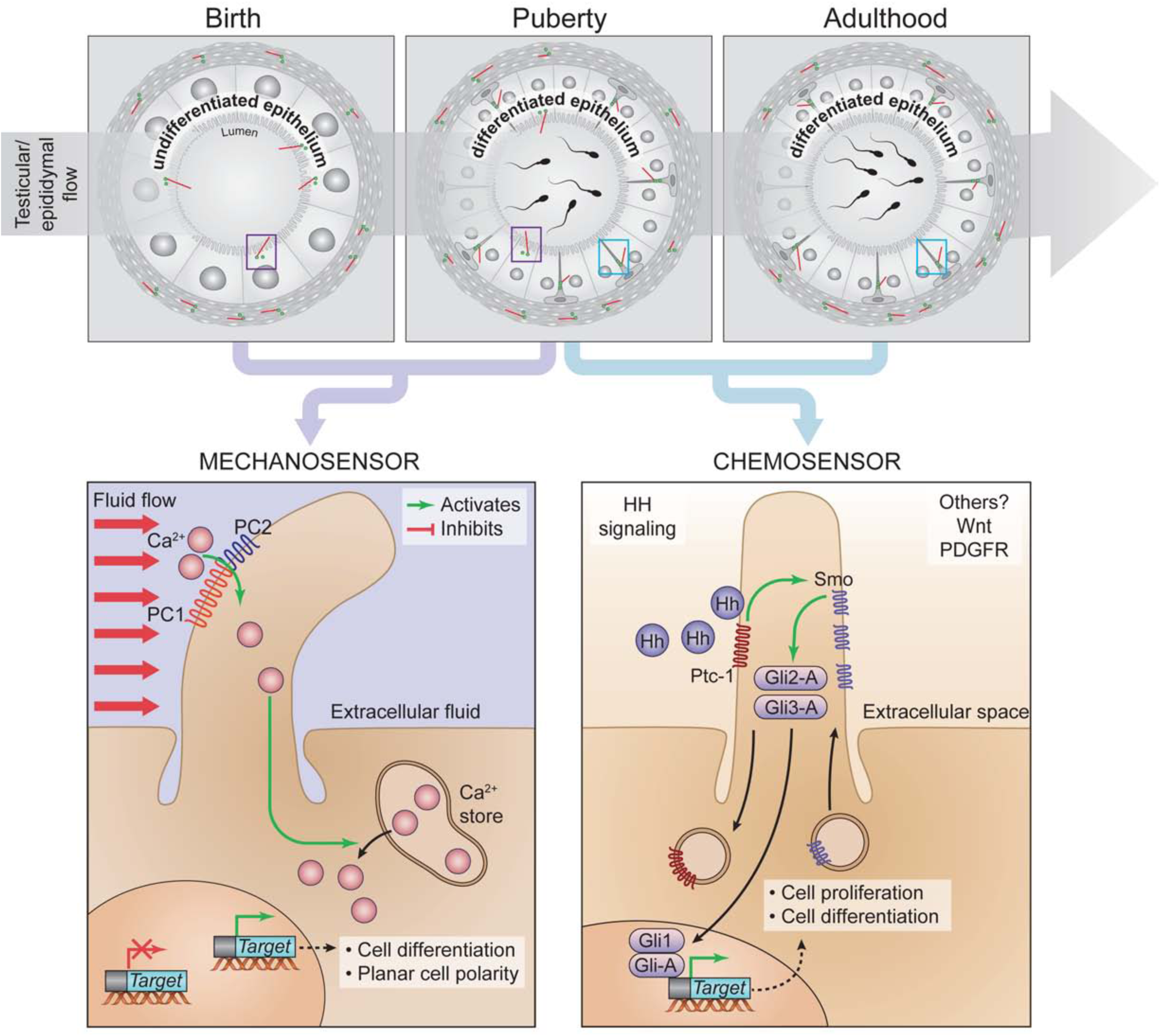
Cell lineage specificity and potential roles of primary cilia in the epididymis during postnatal development. Potential mechano/chemo-sensory functions associated with epithelial cells primary cilia are presented.

By using a transgenic mouse model developed to detect endogenous fluorescence in both ciliary extension and centrioles, we determined that primary cilia were exclusively associated with basal cells in the epididymal epithelium of adult mice. This is consistent with the serendipitous observation made in 2013 by Arrighi, who documented the presence of primary cilia in the epididymis of equine species [18]. In this unique article on epididymis primary cilia, typical 9+0 microtubular patterns and ciliary extensions were observed by transmission electron microscopy associated with basal cells, according to their characteristic morphological features. Our findings not only confirmed the cell specificity of primary cilia though co-localization of primary cilia with keratin-5 positive basal cells, but also underscored a change in ciliary features (length, orientation and signaling components) throughout the different segments of the epididymis. Epididymal basal cells present with dynamic axiopodia extensions that cross the blood-epididymis barrier to sense the lumen [49, 50]. According to our observations, the fact that primary cilia never cross the apical tight junctions in adult epididymis— even when basal cells present with elongated axiopodia—suggests that primary cilia might sense intra-epithelial extracellular fluid rather than the intraluminal fluid in adult mice.

Basal cells from several other stratified and pseudo-stratified epithelia expose primary cilia with distinct signaling functions. For instance, basal cells from the olfactory epithelium have primary cilia that control basal cell activation, proliferation and differentiation, and as such are necessary for regeneration of the epithelium following injury [57]. In addition, primary cilia associated with basal cells from the mammary gland are involved in the acquisition of their stem cell properties via Hh signaling, thereby controlling mammary tissue outgrowth and basal cell tumor development [58]. While limited information is available with regard to basal cell functions in the epididymis, their association with primary cilia could underlie a new functional hypothesis related to epididymis development and homeostasis. According to gene expression profiling, basal cells share common properties with adult stem cells and were recently proposed to differentiate into columnar cells as a mechanism of epididymal epithelium regeneration [59]. Although basal cells from the epididymis do not seem to be progenitors of the other cell populations during early development [14, 60], they may participate in epithelium regeneration via primary cilia signaling, in a similar manner to other epithelia. In addition, primary cilia-dependent control of tumorigenesis via Hh signaling appears to be common in cases of basal cell carcinoma [61]. It is striking that the epididymis rarely develops tumors [62, 63]. Among the few cases that have been described in the literature, two thirds of epididymal epithelial tumours (papillary cystadenoma) are associated with Von Hippel-Lindau (VHL) syndrome, an atypical ciliopathy[64]. Thus, it is possible that primary cilia monitor the epithelium stability of the epididymis and prevent oncogenic events through Hh or Wnt signaling pathways, whose downstream effectors are enriched in basal cells ([59] and our data).

Depending on the physiological context, primary cilia could exert dual sensory functions, either as mechanosensors and/or chemosensors (For reviews, [19, 20, 65]). With regard to chemosensing, the existence of transition fibers located at the base of the cilium physically confines specialized molecules into the cilium, which potentiates signaling responses to extracellular cues. Among these signals, primary cilia coordinate Hh, Wnt and PDGFR and signal-transduction machineries. Interestingly, the components of these pathways—either agonists or downstrean effectors—are all found enriched in basal cells of the epididymis [59] and are particularly relevant to the control of epididymis homeostasis and post-testicular maturation events occurring in this organ [62, 66–71]. For instance, blockage of the Hh signaling pathway following cyclopamine administration triggers a significant decrease in Gli1 and Gli3 transcriptional factor expression as well as a significant decrease in epididymal sperm motility [69]. Whether or not this phenotype relies on primary cilia-dependent Hh signaling remains to be established. Complementary *in vivo* studies on basal cell primary cilia will further define the contribution of this organelle to epididymis physiology, and ultimately to sperm maturation and male fertility.

Finally, the fact that primary cilia are observed at the apical pole of the epithelium from early stages of PND to puberty and colocalize with polycystins (PC1 and PC2), suggests that these organelles might play the role of mechanosensors before puberty. This type of primary cilia-dependent mechanosignaling is broadly documented in the kidney, which shares the same embryonic origin as the epididymis and presents numerous functional similarities with it [24, 26]. In epithelial cells from the distal convoluted and connecting tubules of the kidney, the flow shear stress exerted at the surface of the cells is sensed by primary cilia exposed at the cell surface. The subsequent physical bending of primary cilia triggers PC1 dependent-transepithelial transport of calcium and controls tissue morphogenesis [26]. Approximately 40 years ago, Sun and Flickinger proposed that the flow of testicular-derived fluid entering the epididymis before sperm production may stimulate epithelial cell proliferation and differentiation in the epididymis [1]. Building on our new findings, it is now conceivable that primary cilia are strategically positioned in direct contact with the lumen of the epididymis to initiate shear-flow dependent epithelium maturation and sense the first wave of spermatozoa at puberty to sustain maturation.

In conclusion, our study reveals for the first time the cell lineage specificity of primary cilia along the epididymis at different stages of PND. Acknowledging the important role played by these sensory organelles in most biological systems, our work opens new avenues of research concerning the cellular control of epididymal development and homeostasis. Importantly, since primary cilia components are conserved in the human epididymis, current drugs controlling ciliary functions might constitute new targets for the development of non-hormonal male contraceptives or treatments for cases of unexplained male infertility.

## Acknowledgements

We thank Pr Sylvie Breton, Ph.D., for providing aquaporin 9 antibody used in this project, as well as Pr Marie-Claire Orgebin-Crist, Ph.D., for providing epididymal DC2 cell line. We would like to acknowledge the contribution of Sabine Elowe, Ph.D., for confocal imaging support, Robert Sullivan, Ph.D., for giving us access to the biobank of human tissues, and Christine Légaré, M.Sc., for her technical support. The contributions of Johanne Ouellet in histology services, France Couture in illustration and Julie-Christine Lévesque, M.Sc., from the bio-imagery core facility are also acknowledged. Enthusiastic informal discussions with Rex Ress, Ph.D. were highly appreciated throughout this project.

## Author contributions

Experiments were conceived and designed by ABa, ABe, CB, CRoy, and DS. Contributed to critical reagent, resources and expertise; CB, CRo, DS, JB, MET. Experiments were performed by ABa, ABe, CB, CL, CRoy, DS, OJ, MBR. Data were processed and analyzed by ABa, ABe, CB, CRoy, DS, MBR, and supervised by CB, CRo, DS, JB, and MET. The manuscript was written by ABe, CRoy and CB, and critically reviewed by all authors.

## Supplementary files

**Supplementary file 1.** 3D reconstruction of the epididymal epithelium after ZO-1 staining in adult Arl13b-Cetn2 tg mice

**Supplementary file 2.** Proportion of primary cilia found in basal and myoid cells from different epididymal segments.

**Supplementary file 3.** 3D reconstruction of the epididymal epithelium in Arl13b-Cetn2 tg mice at early stage of PND (at 5 dpn).

**Supplementary file 4.** 3D reconstruction of the epididymal epithelium in Arl13b-Cetn2 tg mice after puberty (at 40 dpn).

**Supplementary file 5.** Immunohistochemistry staining for IFT88 in human epididymis.

